# The MITF-SOX10 regulated long non-coding RNA DIRC3 is a melanoma tumour suppressor

**DOI:** 10.1101/591065

**Authors:** Elizabeth A Coe, Jennifer Y Tan, Michael Shapiro, Pakavarin Louphrasitthiphol, Andrew R Bassett, Ana C Marques, Colin R Goding, Keith W Vance

## Abstract

The MITF and SOX10 transcription factors regulate the expression of genes important for melanoma proliferation, invasion and metastasis. Despite growing evidence of the contribution of long noncoding RNAs (lncRNAs) in cancer, including melanoma, their functions within MITF-SOX10 transcriptional programmes remain poorly investigated. Here we identified 245 candidate melanoma associated lncRNAs whose loci are co-occupied by MITF-SOX10 and that are enriched at active enhancer-like regions. We characterise the function and molecular mechanism of action of one of these lncRNAs, *Disrupted In Renal Carcinoma 3* (*DIRC3*), and show that it operates as a MITF-SOX10 regulated tumour suppressor. *DIRC3* depletion in human melanoma cells leads to increased anchorage-independent growth, a hallmark of malignant transformation, whilst melanoma patients classified by low *DIRC3* expression have decreased survival. *DIRC3* is a nuclear lncRNA that functions locally to activate expression of its neighbouring *IGFBP5* tumour suppressor through modulating chromatin structure and suppressing SOX10 binding to putative regulatory elements within the *DIRC3* locus. In turn, *DIRC3* dependent regulation of *IGFBP5* impacts the expression of genes involved in multiple cancer associated processes. Our work indicates that lncRNA components of the MITF-SOX10 networks are an important new class of melanoma regulators and candidate therapeutic targets.

## INTRODUCTION

Growing evidence supports the biological relevance of gene expression regulation by long non-coding RNAs (lncRNAs) in health and disease. Whilst the molecular functions of most lncRNAs remain poorly understood, investigation of the mode of action of a small number of these transcripts has revealed that they contribute to multiple aspects of gene expression output (1,2). Specifically, a subset of nuclear lncRNAs control the expression of adjacent genes by acting locally, close to their sites of synthesis. Such functions can be mediated by transcript-dependent interactions to facilitate the recruitment of transcription/chromatin regulatory proteins to neighbouring genomic sites and/or to modulate ribonucleoprotein complex assembly (3,4). Alternatively, some nuclear lncRNAs have transcript-independent functions. These can be mediated by the process of RNA polymerase II transcription or splicing, or by the action of DNA-dependent regulatory elements within their loci (5,6). In addition, a small number of lncRNA transcripts have been shown to act in *trans* by moving away from their sites of synthesis to bind and directly regulate multiple target genes on different chromosomes (7–9).

Given their critical regulatory roles, not surprisingly a subset of lncRNAs have been recurrently implicated in cancer. For example, 14.6% of lncRNA loci are found within focal somatic copy number alterations, which contain no cancer associated protein coding genes, in cancer cell genomes (10). LncRNA expression is also dysregulated in most cancers, including melanoma, and lncRNA expression signatures can be used to distinguish different cancers from their normal tissue types (10,11). LncRNAs therefore have potential as cancer biomarkers. A small number of lncRNAs have also been shown to function as oncogenes or tumour suppressors and act as key components of gene regulatory networks controlling carcinogenesis. Such lncRNAs may represent novel therapeutic targets for the development of cancer treatments. For example, activation of the p53 tumour suppressor induces the expression of hundreds of lncRNAs. One of these, *lincRNA-p21* (*TP53COR1*), is needed for p53 dependent repression of anti-apoptotic genes whilst another, *LINC-PINT*, mediates p53-dependent cell cycle arrest (12,13). Similarly, the MYC oncogene binds the promoters of 616 lncRNAs in human B cells, where it directly regulates the expression of some of these (14). Two MYC regulated lncRNAs, *MYCLo-1* and *MYCLo-2*, have also been shown to repress genes that promote colorectal cancer cell proliferation in a MYC-dependent manner (15). These examples illustrate the importance of lncRNA gene regulatory functions downstream of tumour suppressor and oncogenic transcription factors.

Over recent years melanoma has emerged as a leading model for cancer progression, especially with regard to understanding how the microenvironment plays a key role in generating the phenotypic heterogeneity that drives disease progression. It is now apparent that lncRNAs can exert important roles in the biology of melanoma. For example, oncogenic BRAF^V600E^ signalling regulates the expression of 109 putative lncRNAs, one of which *BANCR*, functions as a melanoma oncogene to activate gene expression programmes controlling cell migration (16). The *MIR31HG* lncRNA induces *INK4A*-dependent senescence in response to BRAF^V600E^ expression (17), whereas the lncRNA *SAMMSON* is needed for melanoma proliferation and survival. Targeted inhibition of *SAMMSON* disrupts mitochondrial function and increases the response to MAPK-inhibitor drugs in patient-derived xenograft models of melanoma (18). However, while melanoma associated lncRNAs have great potential for the development of new melanoma treatments, how lncRNAs integrate into the well-defined gene expression networks that underpin different phenotypic states in melanoma is less clear.

The microphthalmia-associated transcription factor MITF plays a critical role in melanocyte development and in melanoma. MITF activates the expression of protein coding genes involved in differentiation and proliferation, DNA replication and repair, mitosis, oxidative phosphorylation and mitochondrial metabolism, and represses the transcription of genes involved in melanoma cell invasion and motility (19–21). *MITF* amplification is frequent in melanoma and particularly common in metastatic forms of the disease where it associates with poor survival (22). *MITF* expression is controlled in part through the action of SOX10, and together MITF-SOX10 co-occupy several thousand binding sites on chromatin to control key gene regulatory networks governing melanocyte development and melanoma (23). Recent work has also reported that MITF-SOX10 drive the proliferative cell state in melanoma and influence the response to MAPKinase inhibiting therapeutics (24,25). Although the SOX10-regulated lncRNA *SAMMSON* is co-amplified with *MITF* in melanoma, our understanding of how MITF-SOX10 drive melanocyte and melanoma biology, and how much of their activity is mediated by lncRNAs is largely unexplored.

Here we identify 245 candidate melanoma associated lncRNAs that are targeted by MITF-SOX10. We show that one of these, *Disrupted In Renal Carcinoma 3 (DIRC3)*, is a clinically important MITF-SOX10 regulated melanoma tumour suppressor gene that controls anchorage-independent growth. Our results reveal that *DIRC3* activates expression of the neighbouring *IGFBP5* tumour suppressor gene to control gene expression programmes involved in cancer. This work highlights that MITF-SOX10 bound lncRNAs, as exemplified by *DIRC3*, can function as important components of MITF-SOX10 expression networks and regulate melanoma growth and progression.

## MATERIAL AND METHODS

### Plasmid construction

To generate pX-dCas9-mod, dCas9 was excised from pAC94-pmax-dCas9VP160-2A-puro (Addgene #48226) as an *AgeI-FseI* fragment and subcloned into pX459 (Addgene #62988) to replace the nuclease active Cas9 protein. A modified sgRNA backbone was then synthesised as a Gblock (IDT) and introduced by cutting the vector with *BbsI* and *BamHI* and replacing the guide backbone with a *BsaI* and *BamHI* digested gene fragment that recapitulated the *BbsI* cloning sites. To generate pX-dCas9-mod-KRAB, the KRAB domain was PCR amplified using pHR-SFFV-dCas9-BFP-KRAB (Addgene #46911) as a template, and cloned into the *FseI* site in the pX-dCas9-mod vector. Five sgRNAs were designed against a genomic region −50 to +150 bp relative to the transcription start site of each target using the Zhang Lab CRISPR Design Tool (https://zlab.bio/guide-design-resources). Linker ligation was performed to insert each sgRNA as a *BbsI* fragment into either pX-dCas9-mod or pX-dCas9-mod-KRAB. We selected two effective sgRNAs for each target and used the published negative control sgRNA sequence described in (26). Oligonucleotides used for cloning are shown in Supplementary Table 1.

### Cell culture and transfections

501mel, SKmel28 and A375 human melanoma cells were cultured in 5% CO^2^ at 37°C in Dulbecco’s modified Eagle’s medium supplemented with 10% fetal bovine serum (growth medium). All cell lines were routinely tested for mycoplasma. For ASO LNA GapmeR (Exiqon) and siRNA (Sigma-Aldrich) mediated knockdown experiments, melanoma cells were seeded in a 6 well plate and transiently transfected using Lipofectamine 2000 reagent (Thermofischer) following the manufacturer’s instructions. 150 pmol each ASO, 100 pmol of siRNA or 1.5 μg MISSION^®^ esiRNA (Sigma-Aldrich) was transfected in each experiment. Experiments to analyse the effect of depletion were carried out 3 days after transfection. For CRIPSRi, 2μg plasmid DNA was transfected per well in a 6 well plate using Lipofectamine 2000 according to the manufacturer’s instructions. Three days later, cells were trypsinised, resuspended in growth medium containing 0.7 μg/ml puromycin and plated onto a 6-cm dish. Drug-resistant cells were grown for 7 days and harvested as a pool. To generate stable cell lines, individual drug resistant clones were isolated and expanded under selection using 1 μg/ml (SKmel28), 1.3 μg/ml (501mel), or 1.5 μg/ml (A375). DIRC3 expression in individual clones was characterised using RT-qPCR. siRNA and ASO sequences are listed in Supplementary Table 1.

### RT-qPCR

RNA was extracted using the GeneJET RNA purification Kit (ThermoFisher Scientific) and reverse transcribed using the QuantiTect Reverse Transcription Kit (Qiagen). Fast SYBR™ Green quantitative PCR was performed on a Step One™ Plus Real-Time PCR System (Applied Biosystems). Sequence of all RT-qPCR primers are shown in Supplementary Table 1.

### Western blotting and cellular fractionation

Western blotting was performed using either anti-MITF antibody rabbit polyclonal (MERCK; HPA003259), anti-SOX-10 (Santa Cruz; sc-365692) or anti-β-ACTIN (Santa Cruz; sc-47778) mouse monoclonal antibodies. For cellular fractionation, approximately 1 × 10^6^ cells were harvested and pelleted at 4 °C. Cell pellet was re-suspended in 200 μl Lysis Buffer (15 mM HEPES, pH 7.5, 10 mM KCl, 5 mM MgCl2, 0.1 mM EDTA pH 8, 0.5 mM EGTA pH 8, 250 mM Sucrose, 0.4% Igepal, 1 mM DTT, 0.5 mM, protease inhibitor cocktail (Roche), 100 U/ml RNAsin) and incubated at 4 °C for 20 minutes with rotation to lyse. Cells were further disrupted by pipetting and centrifuged at 2000 g for 10 minutes at 4 °C. Nuclear (pellet) and cytoplasmic (supernatant) fractions were collected. RNA was then prepared from each fraction and analysed by RT-qPCR.

### Colony forming assays

Approximately 5×10^3^ *DIRC3* depleted or control clonal cells were suspended in 1.5ml of growth medium containing 0.3% noble agar and plated on top of a 0.5% noble base agar layer in a six-well plate. Cells were then grown in 5% CO_2_ at 37°C and supplemented with 100 μl growth medium every three days. After 21 days cells were fixed in 1% formaldehyde and stained using 0.01% crystal violet. Colony number was quantified using ImageJ.

### Transcriptomics

For RNA-sequencing of Hermes melanocytes, IGR37 and IGR39 melanoma cells, cells were grown to 80% confluence and then harvested. Total RNA was purified using the RNeasy mini kit (QIAGEN) according to the manufacturer’s instructions including an on-column DNase digestion step. Library preparation and paired-end sequencing (44-59 million reads per sample) was performed at the Wellcome Trust Centre for Human Genetics, University of Oxford. LncRNA annotation from RNA-seq data was performed as described previously using CGAT pipelines (27,28).

For gene expression analysis, total RNA was prepared in triplicate from knockdown *(DIRC3, IGFBP5)* and control SKmel28 cells using the GeneJET RNA purification Kit (ThermoFisher Scientific). PolyA selected 150-bp paired end RNA sequencing was performed on the Illumina HiSeq4000 (Novogene). A minimum depth of 30M mapped reads were generated per sample. RNA-seq data was analysed as follows. Quality controls were applied for cleaning data for adapters and trimming of low quality sequence ends using trim_galore version 0.4.4. Cleaned data was aligned using FLASH v1.2.11 and mapped to the hg19 genome using bowtie version 1.1.2. The human gene annotation file was obtained from Gencode (v19, genome assembly hg19). Data were aligned to these annotations using the Bioconductor package GenomicAlignments version 1.34.0 function summarizeOverlaps. Statistical analysis was performed with the Bioconductor package DESeq2 (R version 3.5.0, DESeq version 1.22.2). Default settings were used. Differential gene expression was tested between knockdown and control groups. P-values were adjusted by the Benjamini-Hochberg method, controlling for false discovery rate (FDR). Gene Ontology analyses were performed using the Bioconductor package GOstats function hyperGtest. A list of all genes expressed in SKmel28 cells was used as a background dataset. FDR cut-offs were computed from the resulting p-values using the brainwaver package function compute.FDR.

### Gene expression analysis of TCGA melanoma RNA-seq data

Expression correlations were performed as described in (29). Expression data was retrieved using CGDS-R package (https://cran.r-project.org/web/packages/cgdsr/index.html). TCGA samples were ranked according to their expression value of the selected gene (x-axis gene) and plotted as relative expression in black line. The relative *DIRC3* expression (y-axis gene) for each sample was plotted as a bar graph in light grey and the moving average line with a window of 20 was plotted in the same colour as the y-axis. The Spearman correlation coefficient (rho) and an exact P-value of the correlation between the selected genes (x- and y-axis genes) were generated using R function ‘cor.test’.

### Chromatin immunoprecipitation

ChIP was performed as described in (9) using approximately 1×10^7^ SKmel28 clonal CRISPRi cells per assay and 5 μg of the following antibodies: anti-SOX10 rabbit monoclonal antibody (Abcam; ab155279), anti-Histone H3K27ac rabbit polyclonal (Active Motif; #39133) or anti-rabbit IgG control (Merck; PP64). qPCR primers used to amplify SOX10 bound genomic sequences are shown in Supplemental Table 1.

### Chromatin analysis and datasets

H3K4me1, H3K4me3 and H3K27ac ChIP-seq chromatin modification data (GSE58953) from two tumorigenic melanoma cell lines were downloaded from (30). ChIP-seq reads were processed and aligned to NCBI Build 37 (UCSC hg19) as described (30). Reads mapping to lncRNA loci were further estimated using HTseq (version 0.9.1; --minaqual=1, otherwise default parameters; (31)). HA-MITF and SOX10 ChIP-seq (GSE61967) data from (23) were used in this study. NHEK topologically associating domains (TADs) and loop annotated from HiC data were obtained from (32). HiC heatmap was generated using JuiceBox (33). Statistical analyses were performed using the R software environment for statistical computing and graphics (http://www.R-project.org/).

## RESULTS

### Identification of new melanoma-associated lncRNAs targeted by MITF-SOX10

MITF-SOX10-regulated transcriptional programmes play critical roles in the control of melanoma proliferation, invasion and metastasis. Although protein coding gene networks involved in mediating the downstream MITF-SOX10 transcriptional response have been well-defined, the identity and putative functions for MITF-SOX10 regulated lncRNAs in melanoma remain unknown.

To identify candidate lncRNA regulators of melanoma that are part of the MITF-SOX10 network, we used RNA-sequencing (RNA-seq) of Hermes immortalised human melanocytes and two human BRAF^V600E^ mutated melanoma cell lines isolated from the same patient: IGR39 which are MITF-low, de-differentiated and invasive and IGR37 which are MITF-high and non-invasive. This identified a comprehensive set of 11881 intergenic lncRNA transcripts expressed in at least one of these cell types (Fig 1A, Supplementary Table 2). Integration of genome wide maps of HA-tagged-MITF and SOX10 binding in human melanoma from (23) then defined a stringent set of 245 candidate melanoma-associated lncRNAs whose genomic loci are co-occupied by these two transcription factors (Fig. 1A, Supplementary Table 3). Using chromatin state maps from tumorigenic melanocytes (30), we determined that compared to all other melanocyte and melanoma expressed lncRNAs, MITF-SOX10 bound lncRNA loci displayed increased levels of H3K27ac and H3K4me1 modified chromatin, as well as a corresponding higher ratio of the H3K4me1:me3 enhancer-associated chromatin signature, (p<0.05, two-tailed Mann Whitney U test, Fig. 1B and Supplementary Fig. 1). This suggests that MITF-SOX10 bound lncRNAs are enriched at active enhancer-like regions involved in transcriptional control in melanoma. We predict that a subset of these lncRNA transcripts are likely to play an important role in mediating the MITF-SOX10 transcriptional response in melanoma.

**Figure 1.**
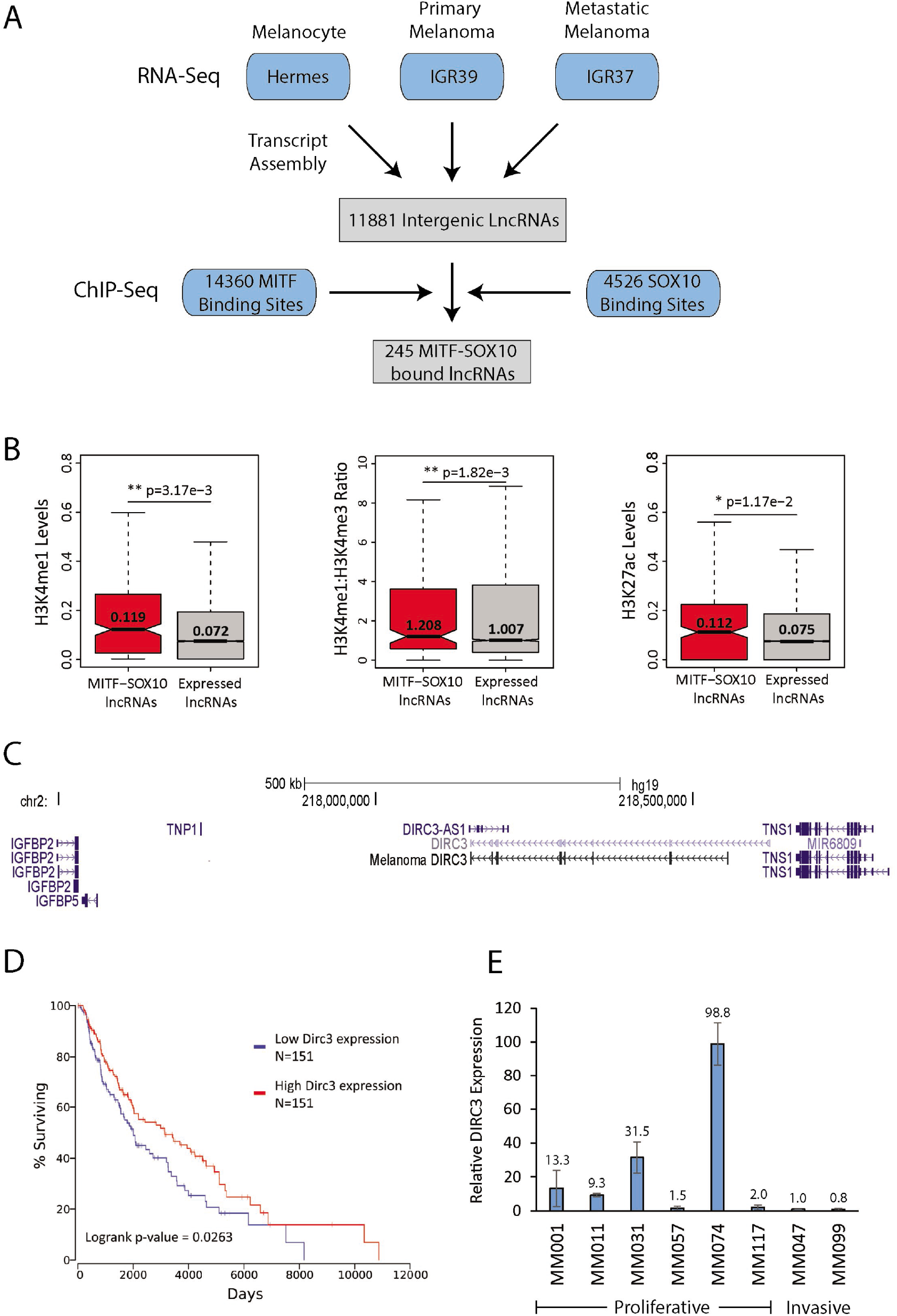
LncRNAs are components of MITF-SOX10 networks in melanoma. (A) Workflow diagram describing the experimental and computational methods used to identify a set of melanocyte and/or melanoma expressed intergenic lncRNAs whose genomic loci are bound by the MITF and SOX10. (B) Distribution of the number of normalised H3K4me1 (left panel), H3K27ac (right panel), and ratio of H3K4me1 to H3K4me3 (middle panel) sequencing reads mapped to MITF-SOX10 bound lncRNAs (red) and all expressed lncRNA (grey) loci in the sh-PTEN HMEL tumorigenic melanoma cell line (30). Differences between groups were tested using a two-tailed Mann-Whitney *U* test, and *p* values are indicated. (C) Schematic illustration of the human *DIRC3* locus and neighbouring protein coding genes (GRCh37/hg19). Rapid amplification of cDNA ends (RACE) and RT-PCR experiments defined *DIRC3* as a 3,384 nucleotide multi-exonic transcript in human melanoma cells. (D) TCGA survival data for Skin Cutaneous Melanoma (SKCM) was linked to *DIRC3* expression using OncoLnc (34). Patients were sorted based on *DIRC3* expression and percent survival compared between *DIRC3* high (top third) and *DIRC3* low (bottom third) groups. Cox regression analysis shows that low *DIRC3* expression correlates with statistically significant decreased survival in melanoma patients (logrank p-value=0.0263). (E) *DIRC3* expression was measured in a panel of proliferative and invasive short term melanoma cultures, as defined in (25), using RT-qPCR. *POLII* was used as a reference gene. Results presented as mean +/- SEM.

To illustrate proof of concept that some MITF-SOX10 bound lncRNAs may comprise an important new class of melanoma regulators we prioritised the *Disrupted In Renal Carcinoma 3* (*DIRC3*) lncRNA, a multi-exonic 3,384 nucleotide transcript in human melanoma cells (Fig. 1C), for further investigation because of the following reasons. (1) Analysis of The Cancer Genome Atlas (TCGA)-derived clinical data using OncoLnc (34) revealed that melanoma patients expressing low levels of *DIRC3* show statistically significant decreased survival compared to those classified based on high *DIRC3* (Fig. 1D). (2) Examination of *DIRC3* adjacent genes suggested that this gene territory encodes important tumour suppressive functions. The human *DIRC3* locus spans approximately 450 kb genomic sequence between the *IGFBP5* and *TNS1* genes (Fig. 1C). Overexpression of the *IGFBP5* mediator of *insulin-like growth factor receptor* (*IGF1R*) signalling inhibits the transformation of human melanoma cells in culture whilst *TNS1* expression is down-regulated in multiple cancers including melanoma (35,36). (3) *DIRC3* expression is higher in proliferative short term melanoma cultures compared to melanomas with an invasive transcriptional signature (Fig. 1E). (4) Evidence for expression of a positionally equivalent *DIRC3* orthologue in mice further indicated that *DIRC3* is a functionally important lncRNA (Supplementary Fig. 2). Together, these data provide strong initial evidence that *DIRC3* may act as a clinically important candidate tumour suppressor gene in melanoma.

### *DIRC3* expression is directly regulated by MITF and SOX10

Analysis of MITF and SOX10 ChIP-seq data in 501mel melanoma cells (23) showed that the *DIRC3* locus contains three sites of MITF-SOX10 co-occupancy: one upstream of the *DIRC3* TSS (BS1) and two within the *DIRC3* gene body (BS3-BS4, Fig. 2A). To test if *DIRC3* is transcriptionally controlled by MITF and SOX10 we used siRNAs to deplete these two transcription factors in 501mel and SKmel28 proliferative-signature human melanoma cell lines, and measured changes in *DIRC3* expression using RT-qPCR. Reduction of MITF mRNA and protein led to increased *DIRC3* expression in both 501mel and SKmel28 cells (Fig. 2B) indicating that *DIRC3* is transcriptionally repressed by MITF in melanoma. In contrast, depletion of SOX10 strongly down-regulated DIRC3 expression in 501mel cells whilst SOX10 knockdown increased *DIRC3* expression in SKmel28 cells (Fig. 2C). This suggests that SOX10 is able to either activate or repress *DIRC3* expression in a cell type-dependent manner.

**Figure 2.**
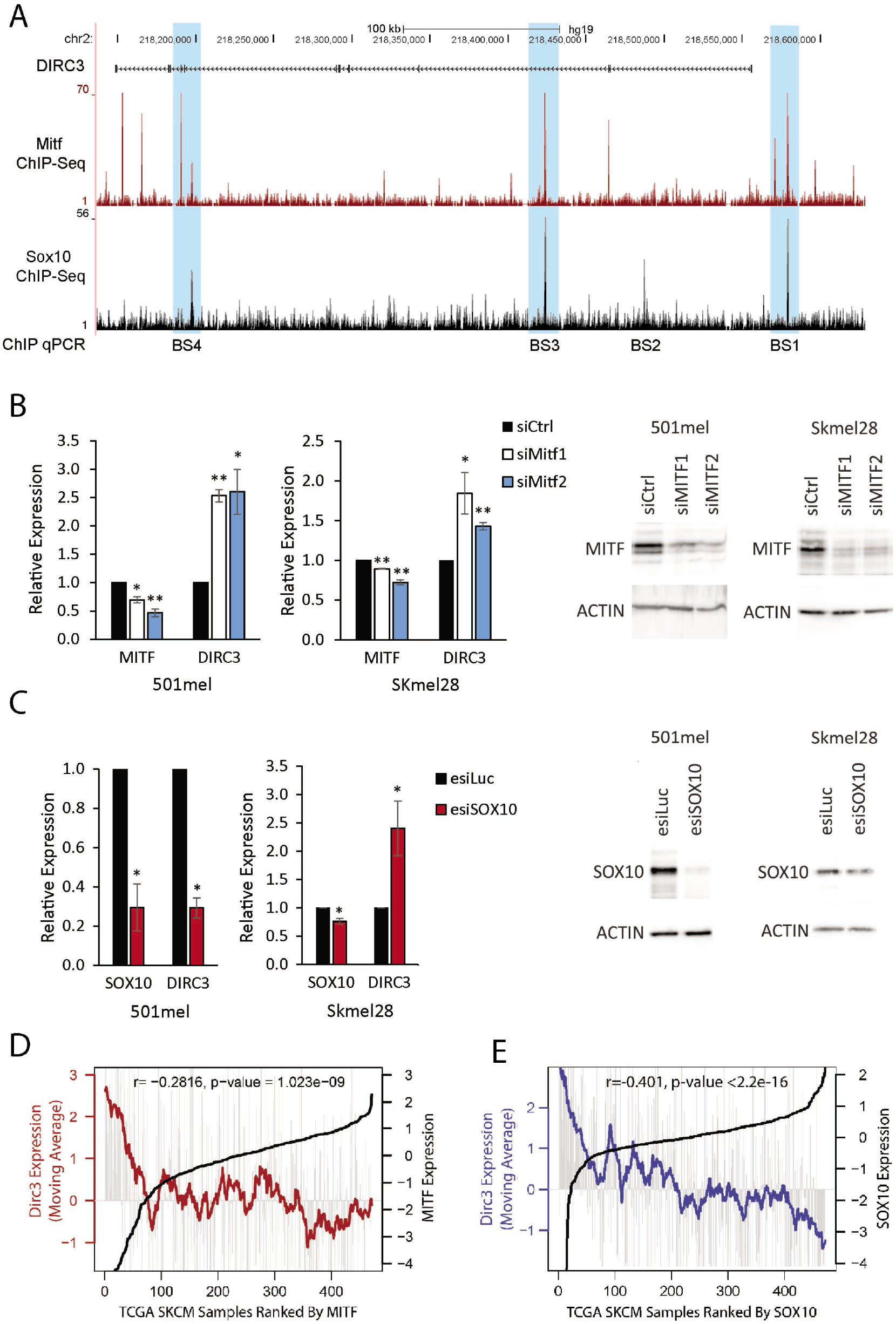
*DIRC3* is a direct MITF and SOX10 transcriptional target. (A) UCSC genome browser view showing that the *DIRC3* locus contains multiple ChIP-seq defined binding sites for MITF and SOX10 in 501mel cells (23). (B, C) MITF represses *DIRC3* (B) whereas SOX10 can both activate and repress *DIRC3* expression (C) in human melanoma cells. MITF and SOX10 were depleted in 501mel and SKmel28 cells using siRNA transfection. Expression changes were analysed using RT-qPCR. *POLII* was used as a reference gene. Results presented as mean +/- SEM., n=3; one-tailed t-test * p<0.05, ** p<0.01. MITF and SOX10 protein levels were analysed by Western blotting. ACTIN was used as a loading control. (D, E) *DIRC3* expression inversely correlates with MITF (D) and SOX10 (E) in melanoma patients. *DIRC3* levels were analysed in 471 TCGA human melanoma samples ranked using increasing MITF or SOX10. Vertical grey lines indicate *DIRC3* expression in each melanoma sample. Moving averages are plotted in bold.

To provide further evidence that DIRC3 is a MITF-SOX10 transcriptional target in melanoma we performed an expression correlation analysis for *DIRC3, MITF* and *SOX10* across the 471 TCGA melanoma RNA-seq samples. This showed that *DIRC3* levels negatively correlate with both MITF and SOX10 in melanoma patients, with *DIRC3* expression being especially high in those tumours with the lowest MITF (Fig. 2D) or SOX10 (Fig. 2E) expression. Taken together, these data provide strong evidence that *DIRC3* is a direct transcriptional target for SOX10 and MITF in melanoma.

### *DIRC3* acts locally to activate expression of the *IGFBP5* tumour suppressor

We next investigated whether *DIRC3* functions to regulate the expression of its adjacent protein-coding genes. Subcellular fractionation experiments first showed that *DIRC3* is a nuclear-enriched transcript in human melanoma cells suggesting that it may function in transcriptional control (Fig. 3A). Control *DANCR* and *NEAT1* lncRNAs were predominantly localised in the cytoplasm and nucleus respectively confirming the efficacy of the fractionation. We then mapped the chromatin structure of the *IGFBP5-DIRC3-TNS1* gene territory using Hi-C data from NHEK cells (32). This showed that *DIRC3* and *IGFBP5*, but not *TNS1*, are located within the same self-interacting topologically associated domain (TAD) and identified two interacting DNA loops between the *DIRC3* locus and the *IGFBP5* gene promoter (Fig. 3B). This indicates that chromatin regulatory interactions bring *DIRC3* into close genomic proximity with *IGFBP5* and suggest a role for *DIRC3* in the local regulation of *IGFBP5*. Consistent with this, expression analysis of TCGA melanoma RNA-seq samples further showed that *DIRC3* levels are positively correlated with *IGFBP5* in melanoma patients (Fig. 3C).

**Figure 3.**
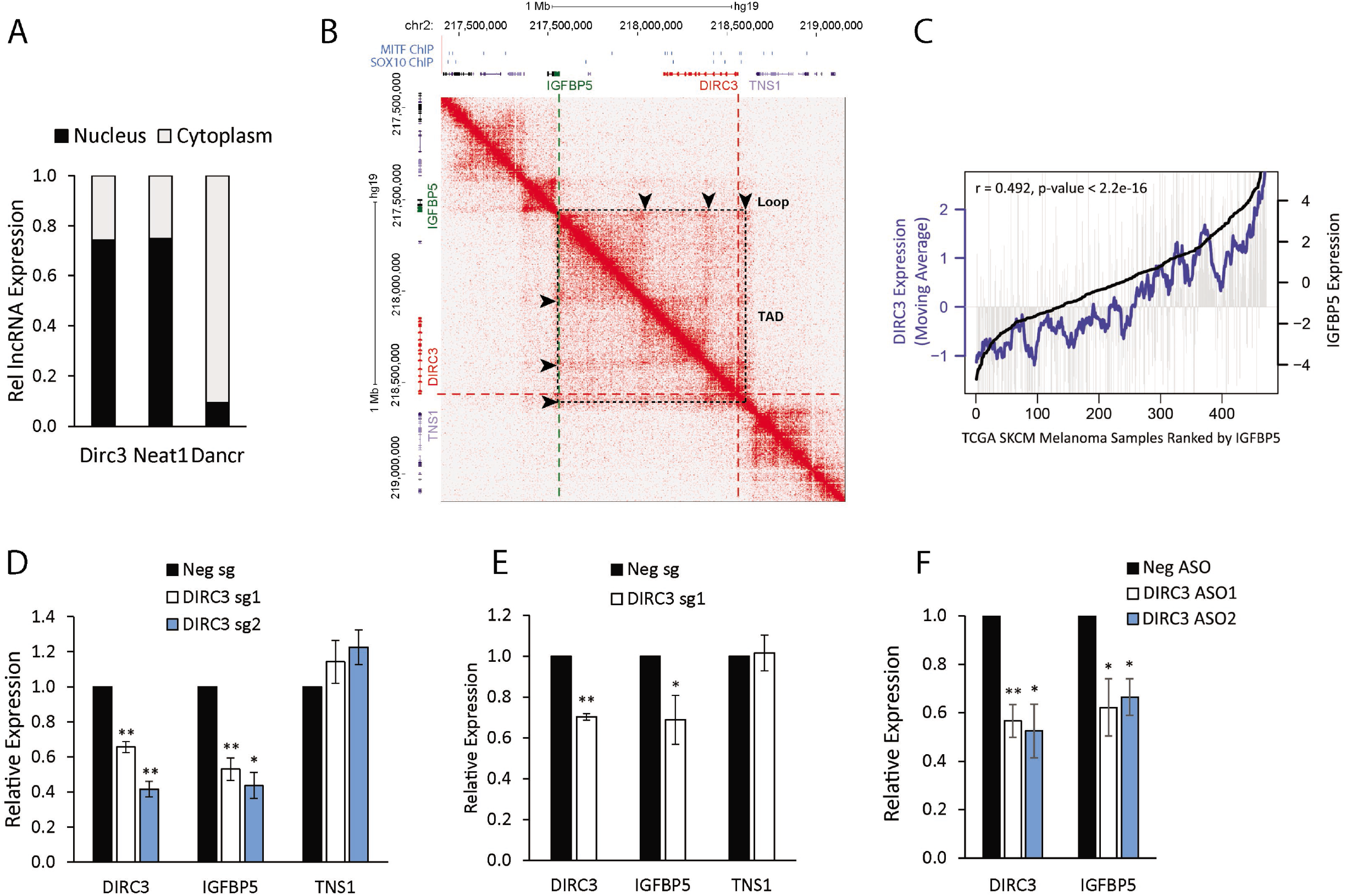
*DIRC3* acts locally to activate expression of the adjacent *IGFBP5* tumour suppressor gene. (A) *DIRC3* transcript is enriched in the nucleus in melanoma cells. SKmel28 cells were biochemically separated into cytoplasmic and nuclear fractions. The relative levels of *DIRC3* and control *DANCR* (cytoplasm) and *NEAT1* (nuclear) transcripts in each fraction were determined by qRT-PCR. (B) *DIRC3* and *IGFBP5* are located within the same TAD. Heatmap displaying chromosomal interactions, measured using HiC, at regions surrounding DIRC3 (red), IGFBP5 (green) and TSN1 (purple), shown in gene browser view, in NHEK (chr2: 217,500,000-219,000,000). The dotted black square box on the heatmap represents a TAD. Chromosomal looping interactions are indicated by black arrows. MITF and SOX10 binding sites are denoted as blue boxes. (C) *DIRC3* expression correlates with *IGFBP5* in melanoma patient samples. Analysis of *DIRC3* expression in TCGA human melanoma samples ranked by increasing *IGFBP5*. Vertical grey lines indicate DIRC3 expression in each melanoma sample. Bold black indicates *IGFBP5* expression; the blue line is the moving average of *DIRC3* expression across 20 melanoma samples. (D-F) *DIRC3* specifically activates *IGFBP5* expression in a transcript dependent manner. DIRC3 levels were depleted by dCas9-KRAB mediated CRISPRi (D), steric hindrance with dCas9 (E) or ASO LNA GapmeR mediated transcript degradation (F) in SKmel28 cells. Expression of *DIRC3* and the indicated neighbouring genes were measured using RT-qPCR with results normalised to *POLII*. Mean values +/- SEM, n = 3. One-tailed student’s t-test p < 0.05 * p < 0.01 **.

We therefore tested whether *DIRC3* transcriptionally regulates *IGFBP5*. To do this, CRISPR interference (CRISPRi) was performed to deplete *DIRC3* expression in Skmel28 cells using two different sgRNAs *(DIRC3sg1* and *DIRC3sg2)* to target the catalytically inactive dCas9-KRAB transcriptional repressor to the *DIRC3* promoter. RT-qPCR analysis of neighbouring gene expression showed that *DIRC3* depletion specifically down-regulated expression of the 3’ IGFBP5 gene, when compared to a control non-targeting guide, whilst the levels of the 5’ *TNS1* gene did not change (Fig. 3D). To validate the specificity of this regulation we also used the *DIRC3sg1* RNA to precisely target dCas9 alone to the *DIRC3* TSS and block *DIRC3* transcription by steric hindrance of RNA polymerase recruitment. RT-qPCR confirmed that *DIRC3* down-regulation also reduced *IGFBP5* expression using this approach whilst *TNS1* levels did not change (Fig. 3E). LncRNA gene regulatory functions can be mediated by the RNA molecule or by the process of RNA polymerase II transcription and/or splicing (2). To test whether *DIRC3* function was transcript-dependent, we used locked nucleic acid (LNA)-modified anti-sense oligonucleotides (ASOs) to deplete *DIRC3* transcript levels and measured changes in *IGFBP5* in SKmel28 cells. This showed that an approximately 50% reduction in the levels of *DIRC3* using two different ASOs resulted in a specific down-regulation of *IGFBP5* compared to a non-targeting control ASO (Fig. 3F).

Together, these data indicate that *DIRC3* is a nuclear localised transcriptional regulator that acts in a transcript-dependent manner to activate expression of the adjacent *IGFBP5* tumour suppressor gene in melanoma.

### *DIRC3* regulates IGFBP5-dependent gene expression programmes involved in cancer

We next performed RNA-seq of *DIRC3* knockdown SKmel28 cells to define the genome-wide impact of *DIRC3* expression in melanoma. ASO-mediated depletion of *DIRC3* expression by approximately 65% resulted in significant changes in the expression of 1886 genes (at a 5% false discovery rate (FDR)) compared to a non-targeting control (Fig. 4A and Supplementary Table 4). 1015 (54%) of these genes were up-regulated and 871 (46%) were down-regulated upon *DIRC3* loss of function. To determine the extent by which *DIRC3* transcriptional response is mediated through *IGFBP5* regulation, we also performed RNA-seq of SKmel28 cells in which *IGFBP5* expression was reduced by approximately 80%. This identified 557 differentially expressed genes (5% FDR) compared to a negative control (Fig. 4B and Supplementary Table 5). We then analysed the intersection of *DIRC3* and *IGFBP5* regulated genes and identified 240 common targets (Fig. 4C and Supplementary Table 6). This overlap represents a significant 22.8-fold enrichment (p < 1e-6) over the expected number based on random sampling of all expressed genes. Moreover, the expression levels of the majority of shared target genes change in the same direction following either *DIRC3* or *IGFBP5* depletion, providing further confirmation that *DIRC3* up-regulates *IGFBP5* (Fig. 4D). Gene ontology (GO) enrichment analysis revealed that *DIRC3-IGFBP5* shared targets are significantly enriched for regulators of cell migration, proliferation, differentiation and metabolism (Fig. 4E and Supplementary Table 7). *DIRC3* therefore possesses *IGFBP5*–dependent gene regulatory functions in melanoma that act to control cancer associated biological processes.

**Figure 4.**
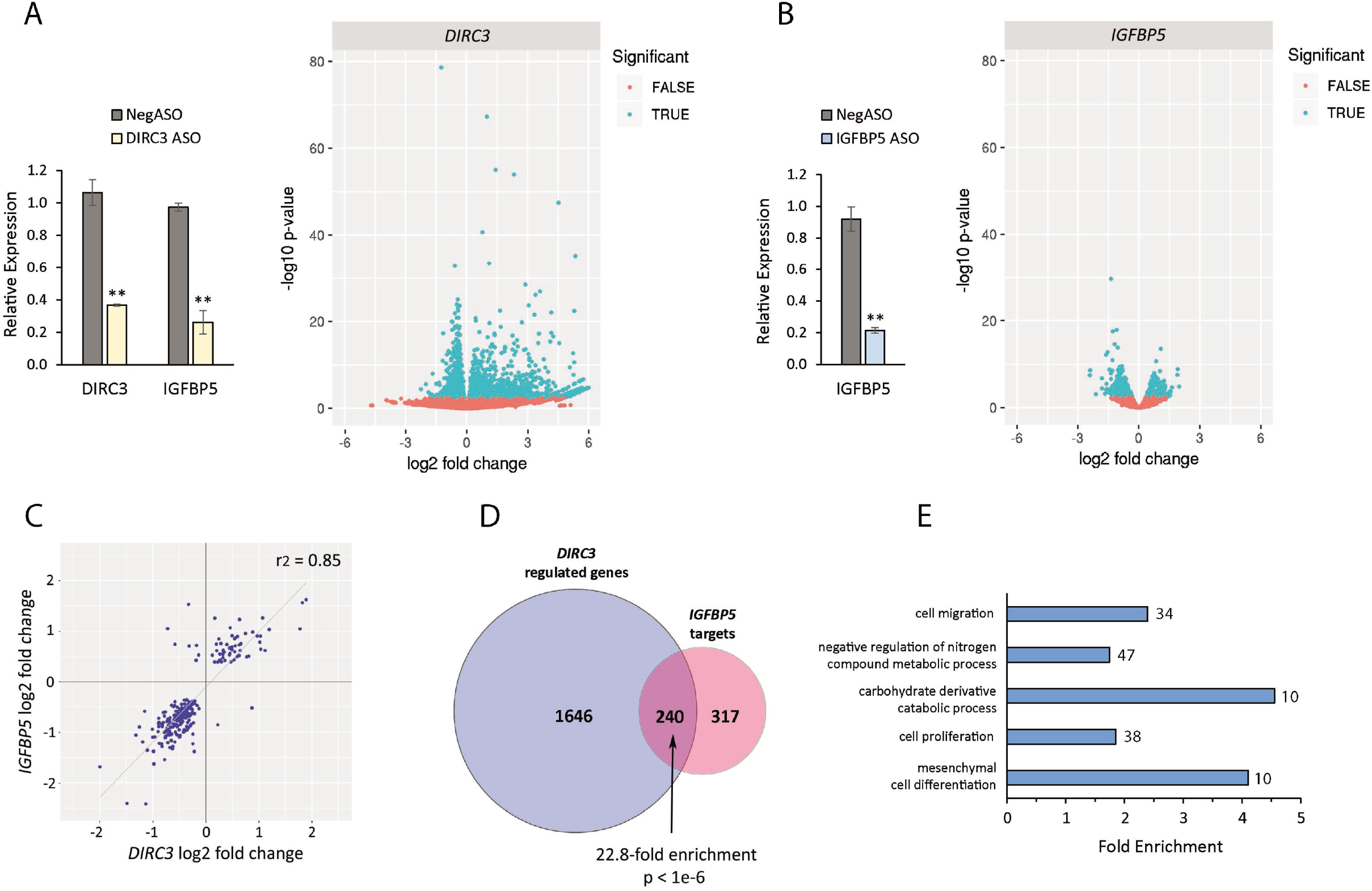
*DIRC3* regulates IGFBP5-dependent gene expression programmes involved in cancer. (A) *DIRC3* depletion induces statistically significant changes (FRD 5%) in the expression of1886 genes in SKmel28 cells using RNA-seq. (B) RNA-seq of *IGFBP5* knockdown SKmel28 cells identifies 557 IGFBP5 target genes (FDR 5%). (C) Intersection of *DIRC3*- and IGFBP5-regulated genes detects 240 common targets. (D) The expression levels of all *DIRC3-IGFBP5* shared target genes change in the same direction following either *DIRC3* or *IGFBP5* depletion. (E) Gene Ontology enrichment analysis of *DIRC3-* and *IGFBP5* shared target genes was performed using GOstats and FDR correction was applied. Representative significantly enriched categories are shown and the number of genes found in each category are indicated.

### *DIRC3* loss of function increases anchorage-independent growth of multiple melanoma cell lines

We next used CRISPRi to generate *DIRC3* loss-of-function cell lines to investigate the role of *DIRC3* in controlling cellular transformation in melanoma. To do this, *DIRC3* expression was first depleted in SKmel28 cells by stable transfection of a plasmid co-expressing the dCas9-KRAB transcriptional repressor along with either the *DIRC3* targeting sgRNAs (*DIRC3sg1* and *DIRC3sg2*) or a negative control. We generated two independent clonal lines, using distinct sgRNAs, in which *DIRC3* expression was depleted by (~70%) compared to a negative control (Fig. 5A, left panel), and showed using RT-qPCR these lines also displayed reduced *IGFBP5* (Fig. 5A, middle panel). *DIRC3* depleted SKmel28 cells proliferate to similar levels as control (Supplementary Fig. 3). As anchorage-independent growth is a good predictor of melanoma metastasis *in vivo* (37), we thus assayed the effect of *DIRC3* loss-of-function on growth in soft agar. We found that stable reduction of *DIRC3* expression, using different dCas9-KRAB targeting sgRNAs, led to a significant increase in melanoma cell colony formation in soft agar (Fig 5A, right panel).

**Figure 5.**
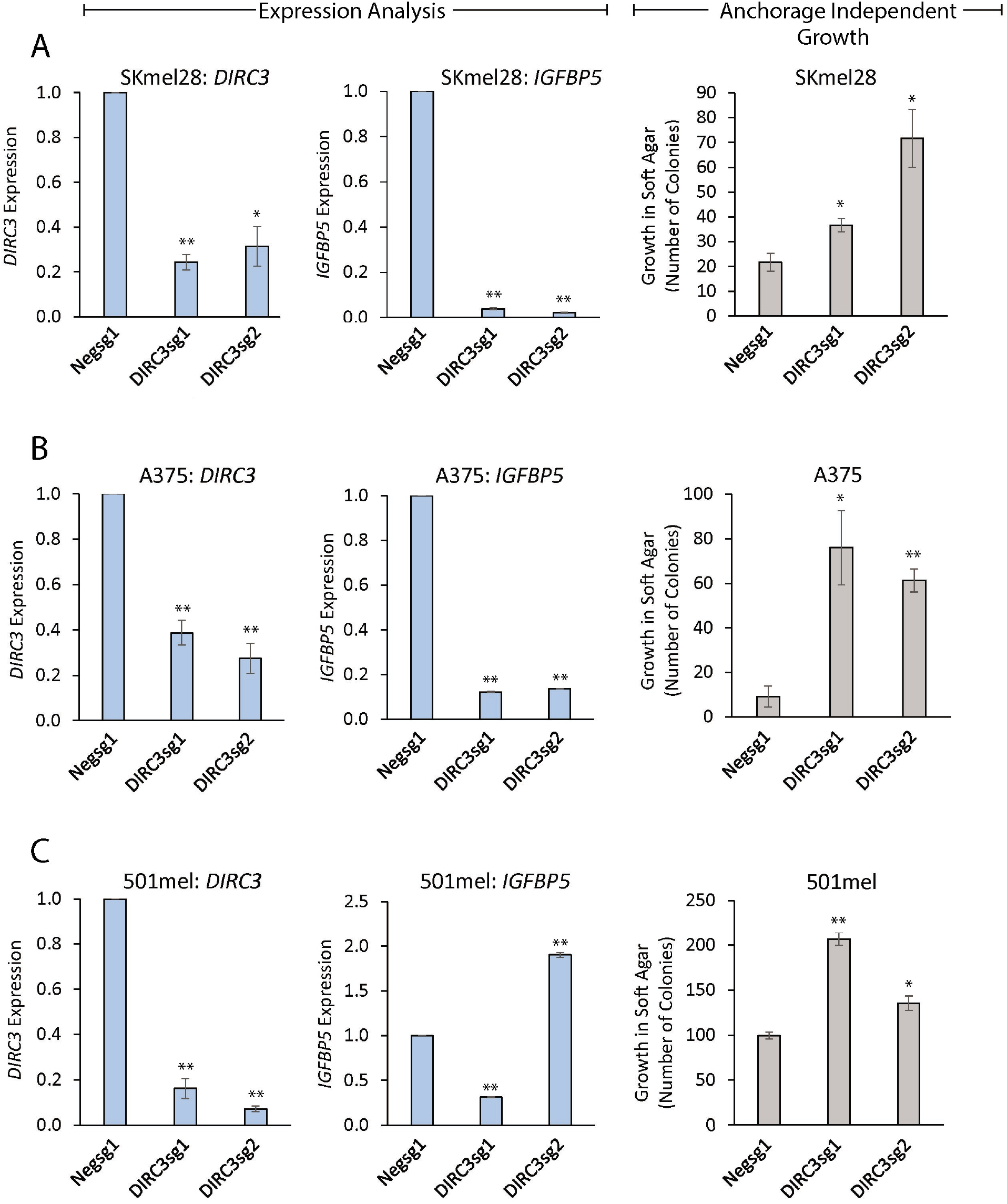
*DIRC3* depletion increases the anchorage-independent growth of melanoma cells in soft agar. *DIRC3* expression was stably depleted in SKmel28 (A), A375 (B) and 501mel (C) human melanoma cells using CRISPRi with two independent DIRC3 targeting sgRNAs and a non-targeting negative control. *DIRC3* (left panel) and *IGFBP5* (middle panel) levels were determined using RT-qPCR. Results are presented as mean values +/- SEM, n = 4. One-tailed student’s t-test p < 0.05 * p < 0.01 **. *DIRC3* depleted and control cell lines were seeded on soft agar and colony formation quantified 21 days later using ImageJ. Mean values +/- SEM, n = 4 (right panel).

We then assessed the generality of DIRC3-mediated control of anchorage-independent growth in melanoma and extended our loss of function analysis to include additional *DIRC3* expressing melanoma cell lines. The results showed that CRISPRi mediated stable depletion of *DIRC3* in A375 and 501mel melanoma cells, using either *DIRC3sg1* or *DIRC3sg2* to target dCas9-KRAB to the *DIRC3* promoter, also resulted in significantly increased colony formation in soft agar compared to control (Fig. 5B, C). This indicates that *DIRC3* regulation of anchorage-independent growth in melanoma is cell line-independent and is in agreement with TCGA clinical data (Fig. 1D) showing that melanoma patients expressing low levels of *DIRC3* display decreased survival, compared to those classified based on high *DIRC3* expression. Additionally, RT-qPCR analysis revealed that *IGFBP5* expression was suppressed in 6 out of 7 of *DIRC3* loss of function cell lines and that the cell line in which *IGFBP5* was no longer depleted showed the smallest increase in colony number (Fig. 5, middle and right panels). These results provide further evidence that *DIRC3* acts through *IGFBP5* to regulate growth in soft agar. Taken together, these results define *DIRC3* as a new melanoma tumour suppressor gene that acts to inhibit the anchorage-independent growth of melanoma cells in culture.

### *DIRC3* regulates local chromatin structure to block SOX10 DNA binding and activate *IGFBP5*

MITF-SOX10 co-occupancy marks active regulatory elements within transcriptional enhancers in melanoma cells (23). The *DIRC3* locus overlaps multiple MITF-SOX10 bound sequences. *DIRC3* thus represents an exemplar to test whether MITF-SOX10 bound lncRNAs can modulate the downstream MITF-SOX10 transcriptional response in melanoma by regulating the association of these transcription factors with their binding sites in chromatin. To do this, we performed ChIP-qPCR in control and *DIRC3* depleted cells and used SOX10 to determine changes in transcription factor occupancy at its binding sites within the *DIRC3* locus. We were unable to identify an anti-MITF antibody to work effectively in ChIP experiments. We first confirmed SOX10 binding at the sites mapped by ChIP-seq within the *DIRC3* locus and at the *DIRC3* promoter in SKmel28 cells. We then found that following CRISPRi mediated stable depletion of *DIRC3*, using either *DIRC3*sg1 or *DIRC3*sg2 dCas9-KRAB targeting sgRNAs, SOX10 protein levels were not affected (Fig. 6A), but that SOX10 chromatin occupancy was significantly increased at its binding sites (Fig. 6B). *DIRC3* therefore functions locally at its site of synthesis to block SOX10 binding at putative regulatory elements within its locus.

**Figure 6.**
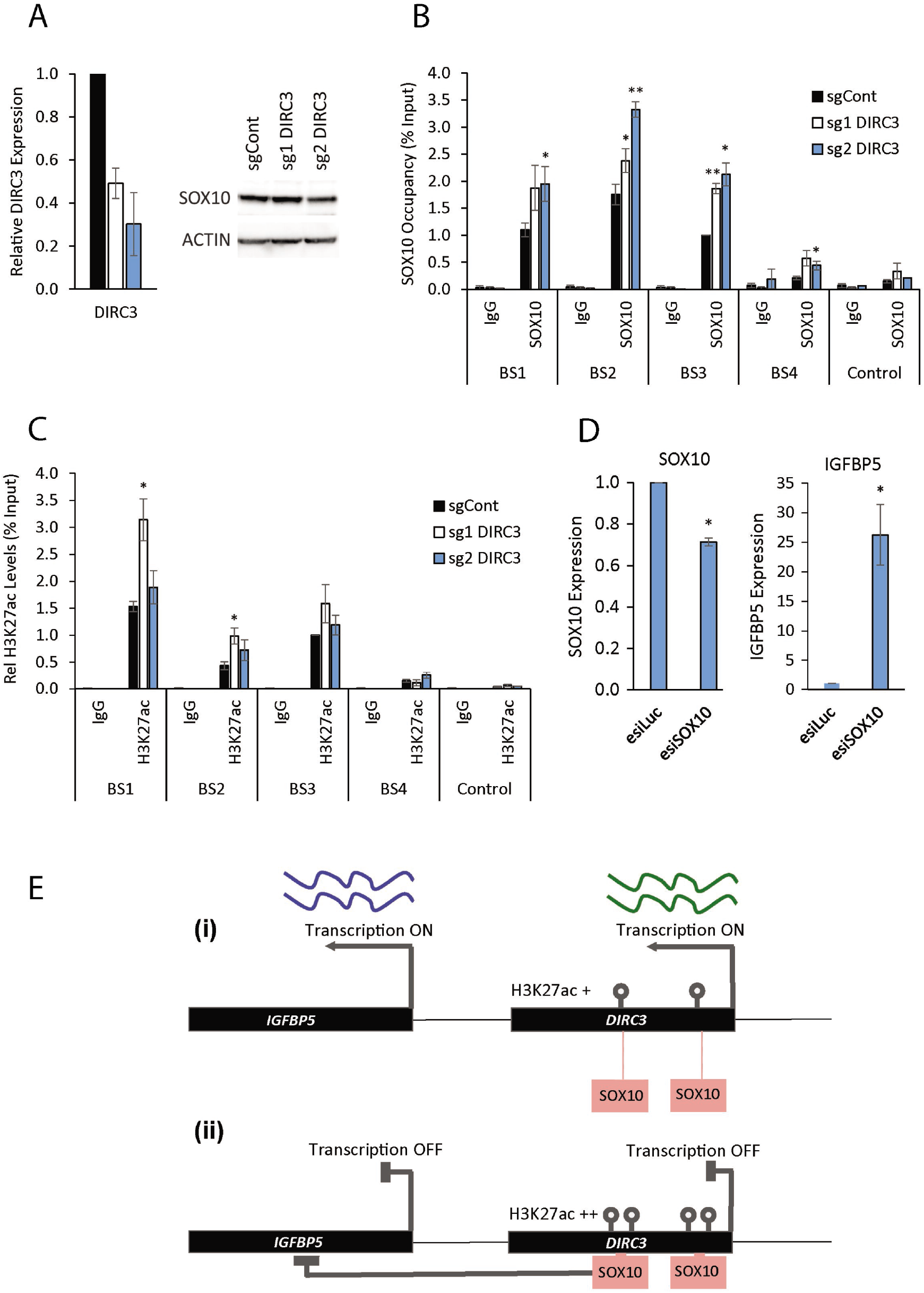
*DIRC3* induces closed chromatin at its site of expression thereby blocking SOX10 DNA binding and activating *IGFBP5*. ChIP assays were performed in *DIRC3* depleted SKmel28 and control cell lines using the indicated antibodies against either SOX10, H3K27ac or an isotype specific IgG control. (A) *DIRC3* depletion was confirmed using qRT-PCR. Western blotting showed that SOX10 protein levels do not change upon *DIRC3* knockdown. ACTIN was used as a loading control. (B) The indicated SOX10 binding sites were analysed by qPCR. % input was calculated as 100*2^(Ct Input-Ct IP). (C) DIRC3 depletion leads to an increase in H3K27ac levels at SOX10 bound regulatory elements within the *DIRC3* locus. (D) SOX10 represses IGFBP5 expression. *SOX10* levels were reduced in SKmel28 cells using esiRNA transfection. *SOX10* and *IGFBP5* expression was quantified using RT-qPCR three days later. *POLII* was used as a reference gene. (E) Model illustrating that *DIRC3* acts locally to close chromatin and block SOX10 chromatin binding at melanoma regulatory elements within its locus. This leads to a block in SOX10 mediated repression of *IGFBP5* and subsequent increase in *IGFBP5* expression. All qPCR results are presented as mean values +/- SEM, n = 3. One-tailed student’s t-test p < 0.05 * p < 0.01 **.

Active transcriptional regulatory elements are marked by H3K27ac modified chromatin. ChIP-qPCR showed that the SOX10-bound sites at *DIRC3* are specifically enriched for H3K27ac when compared to both an isotype control antibody and non-bound regions within the *DIRC3* locus (Fig. 6C). These locations are thus likely to represent DNA sequences involved in transcriptional control in melanoma. Furthermore, stable depletion of *DIRC3* using CRISPRi led to a significant increase in H3K27ac at SOX10 bound sequences within *DIRC3* in SKmel28 cells using ChIP-qPCR (Fig. 6C). This suggests that *DIRC3* functions at its site of expression to close local chromatin structure and control transcription factor occupancy.

We then investigated whether *IGFBP5* is a SOX10 transcriptional target and showed using RT-qPCR that SOX10 depletion increased *IGFBP5* expression approximately 25-fold in SKmel28 cells (Fig. 6D). These data suggest that SOX10 directly represses *IGFBP5* expression in melanoma. Our results are therefore consistent with a model in which *DIRC3* acts locally to block SOX10 chromatin binding at melanoma regulatory elements and activate *IGFBP5* expression (Fig. 6E). We propose that MITF-SOX10 bound lncRNAs, such as *DIRC3*, which function using this mechanism of action have potential to modify the MITF-SOX10 transcriptional response in melanoma.

## DISCUSSION

MITF and SOX10 co-occupy approximately 3,600 binding sites on chromatin in human melanoma cells, marking a subset of active regulatory elements that function to control key melanoma transcriptional programmes underpinning proliferation, invasion and metastasis (23). Previous studies have largely focussed on the ability of MITF-SOX10 to exert their effect on melanoma biology via regulation of protein-coding genes. Here we show that lncRNAs are also important components of MITF-SOX10-driven transcriptional programmes and identify 245 candidate melanoma-associated lncRNAs whose loci are co-bound by MITF and SOX10. These genes are marked by an increased ratio of H3K4me1:H3K4me3 modified chromatin, when compared to all melanocyte lineage lncRNAs identified in our study, suggesting that they preferentially overlap active enhancer-like transcriptional regulatory elements important for melanoma.

To exemplify the importance of MITF-SOX10-regulated lncRNAs in melanoma biology, we prioritised one, *DIRC3*. Significantly we found that *DIRC3* loss of function in three melanoma cell lines leads to increased anchorage-independent growth, a strong predictor of the metastatic potential of melanoma cells (37). Mechanistically, we reveal that *DIRC3* is a nuclear regulatory lncRNA that functions in a transcript-dependent manner to activate expression of its adjacent gene, *IGFBP5*. We show that *IGFBP5* and *DIRC3* are located within the same self-interacting TAD in chromatin and demonstrate that *DIRC3* depletion leads to increased SOX10 occupancy and H3K27ac at putative regulatory elements within the *DIRC3* locus. This results in a concomitant increase in SOX10 mediated repression of *IGFBP5*. These data suggest that *DIRC3* acts at its site of synthesis to modify chromatin structure and control *IGFBP5* regulatory element activity in melanoma. This is important as dysregulated *IGFBP5* transcriptional control can act as a driver of cancer growth and metastasis, regulating cell proliferation, differentiation and metabolism using both IGF1R-dependent as well as - independent mechanisms of action (38,39). For example, copy number alterations encompassing an *IGFBP5* enhancer on 2q35 modulate breast cancer risk (40). In melanoma, a recent preprint reported that inactivation of *IGFBP5* distal enhancers down-regulates *IGFBP5* expression and promotes melanomagenesis by inducing an *IGF1R-AKT* signalling-dependent increase in glycolysis and metabolic reprogramming (41). Consistent with this, *IGFBP5* negatively regulates IGFR1 and MAPK kinase signalling to inhibit melanoma proliferation and metastasis, whilst *IGFBP5* overexpression reduces the anchorage-independent growth of A375 melanoma cells (36). Given the key role of *IGFBP5* in melanoma biology, its regulation by *DIRC3* downstream from MITF-SOX10 is likely significant. Indeed, transcriptomic analysis of *DIRC3* and *IGFBP5* loss of function cells showed that *DIRC3* and *IGFBP5* common target genes are enriched for regulators of cancer associated processes such as cell migration, proliferation and metabolism, as well as mesenchymal cell differentiation. Our results therefore suggest that *DIRC3* acts through *IGFBP5* to exert its tumour suppressive effect in melanoma.

The ability of melanoma cells to switch between proliferative and invasive cell states is important for subpopulations of cells within a heterogeneous tumour to gain invasive properties and metastasize (25,42). Our finding that *DIRC3* expression is lower in invasive compared to proliferative melanomas suggests that down-regulation of *DIRC3* may be an important event in the development of melanoma. This is consistent both with its role as a tumour suppressor and with the observation that melanoma patients with low levels of *DIRC3* display decreased survival. Previous studies have shown that high MITF-SOX10 expression defines the proliferative cell state in melanoma and that switching between different states is accompanied by large scale changes in chromatin structure (25,42). Our data suggests that MITF-SOX10 bound lncRNAs, as exemplified by *DIRC3*, may regulate chromatin accessibility at their sites of expression. Furthermore, the discovery that *DIRC3* blocks SOX10 binding to putative regulatory regions within its locus suggests that lncRNA components of MITF-SOX10 networks can act not only as downstream mediators of MITF-SOX10 function but as feedback regulators of MITF-SOX10 transcriptional activity. Such melanoma associated lncRNAs thus have potential to play widespread roles in fine tuning the MITF-SOX10 transcriptional response and may act as important regulators of cell state transitions in melanoma. In particular, our data indicates that *DIRC3* is a novel and clinically important melanoma tumour suppressor gene. We predict that driving up-regulation of *DIRC3* expression may represent a new therapeutic strategy for melanoma.

## ACCESSION NUMBERS

All RNA-seq data will be deposited in the GEO database.

## SUPPLEMENTARY DATA

Supplementary Data are available online.

## ACKNOWLEDGEMENT

We thank Dr Lesheng Kong, Dr Michael Clark and Prof Chris Ponting for help with bioinformatics; Dr Robert Siddaway for sharing HA-MITF ChIP-seq data; and Dr Eleonora Leucci for kindly providing RNA samples from short term melanoma cultures.

