## Supplementary material for "The MITF-SOX10 regulated long non-coding RNA DIRC3 is a melanoma tumour suppressor": Supplental Figures

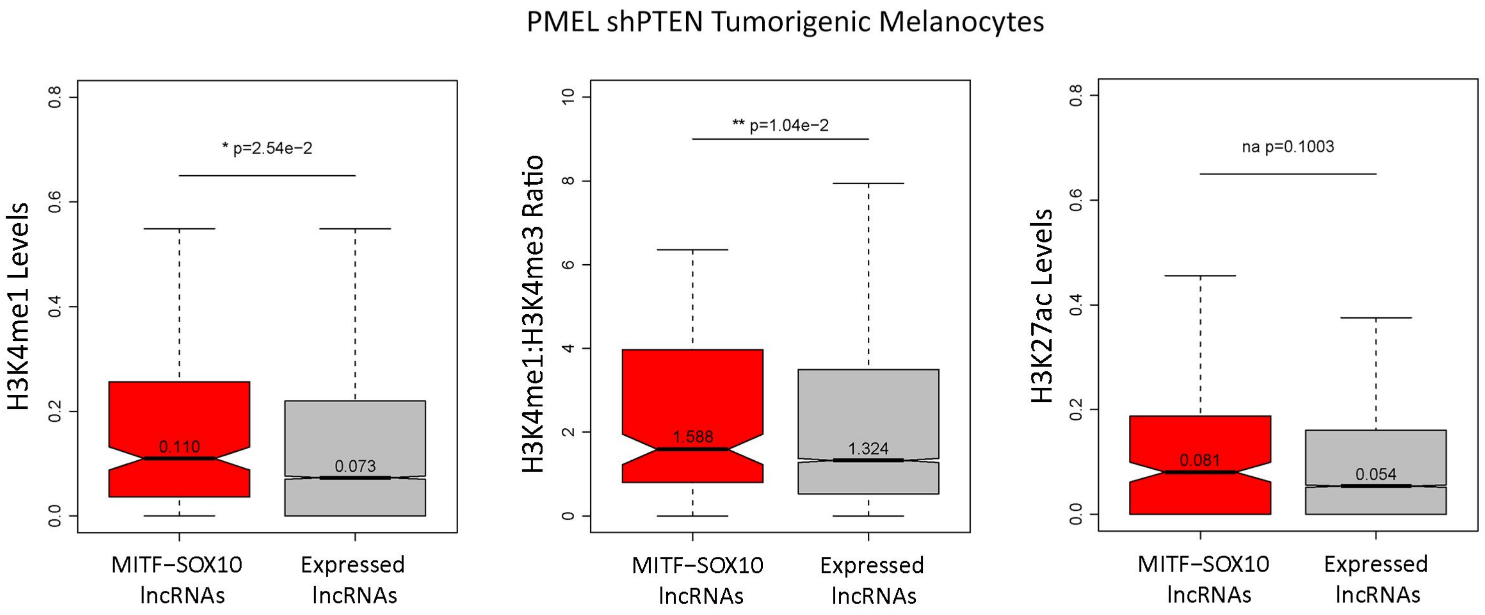


Figure S1 MITF-SOX10 bound lncRNAs are enriched at enhancer-like regions. Distribution of the number of normalised H3K4me1 (left panel), H3K27ac (right panel), and H3K4me1:H3K4me3 ratio (middle panel) sequencing reads mapped to MITF-SOX10 bound lncRNAs (red) and all expressed lncRNA (grey) loci in an additional tumorigenic melanoma cell line (sh-PTEN PMEL cells). Differences between groups were tested using a two-tailed Mann-Whitney U test, and p values are indicated.

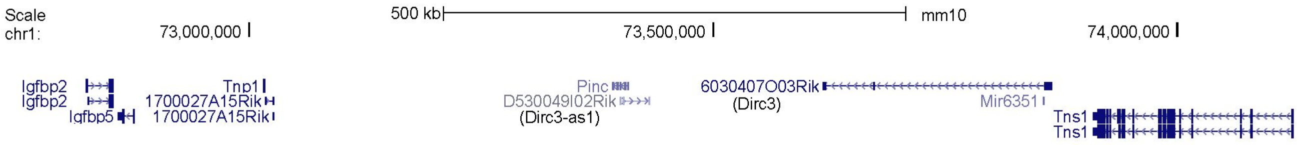


Figure S2. Schematic illustration of the positionally equivalent mouse Dirc3 and neighbouring protein coding genes (GRCm38/mm10).

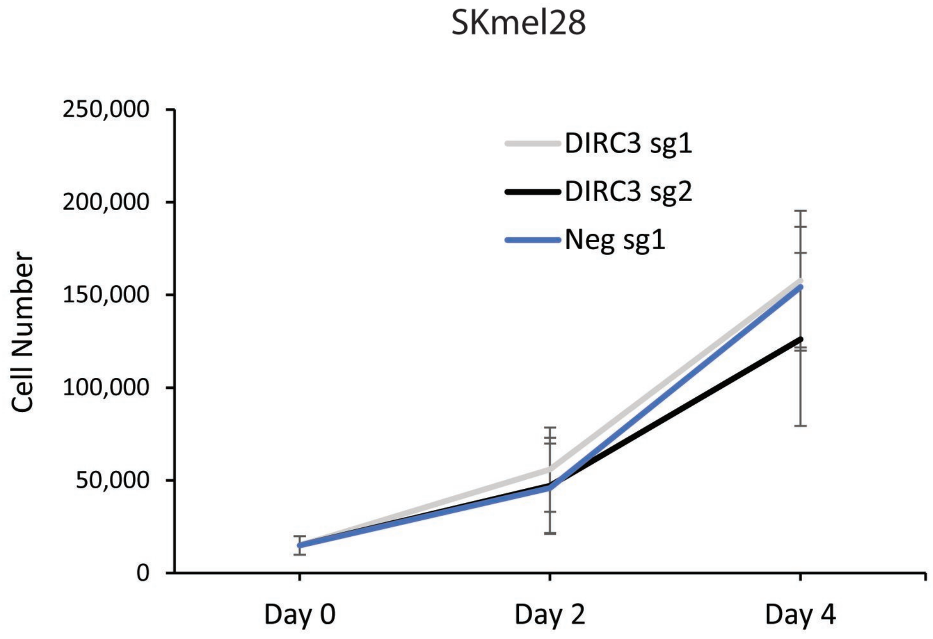


Figure S3. *DIRC3* depleted SKmel28 cells proliferate to similar levels as control. *DIRC3* CRISPRi and control clonal knockdown SKmel28 cells were seeded at a density of 1500 cells per well in a 6-well plate and grown at $37^0$C in 5% $CO^2$. The number of cells were counted 2 and 4 days. n = 3. Mean values +/- SEM.
